# A family of transcription factors that limit lifespan: ETS factors have conserved roles in longevity

**DOI:** 10.1101/438879

**Authors:** Adam J. Dobson, Richard Boulton-McDonald, Lara Houchou, Ziyu Ren, Mimoza Hoti, Maria Rodriguez-Lopez, Alexis Gkantiragas, Afroditi Gregoriou, Jürg Bähler, Marina Ezcurra, Nazif Alic

**Affiliations:** Institute of Healthy Ageing, Department of Genetics, Evolution and Environment, University College London, Gower Street, London WC1E 6BT, UK; School of Biological and Chemical Sciences, Queen Mary University of London, London E1 4NS, UK

## Abstract

Increasing average population age, and the accompanying burden of ill health, is one of the public health crises of our time. Understanding the basic biology of the ageing process may help ameliorate the pathologies that characterise old age. Ageing can be modulated, often through changes in gene expression where regulation of transcription plays a pivotal role. Activities of Forkhead transcription factors (TFs) are known to extend lifespan, but detailed knowledge of the broader transcriptional networks that promote longevity is lacking. This study focuses on the E twenty-six (ETS) family of TFs. This family of TFs is large, conserved across metazoa, and known to play roles in development and cancer, but the role of its members in ageing has not been studied extensively. In *Drosophila*, an ETS transcriptional repressor, *Aop*, and an ETS transcriptional activator, *Pnt*, are known to genetically interact with *Foxo* and activating *Aop* is sufficient to extend lifespan. Here, it is shown that *Aop* and *Foxo* effect a related gene-expression programme. Additionally, *Aop* can modulate *Foxo*’s transcriptional output to moderate or synergise with *Foxo* activity depending on promoter context, both *in vitro* and *in vivo*. *In vivo* genome-wide mRNA expression analysis in response to *Aop*, *Pnt* or *Foxo* indicated, and further experiments confirmed, that combinatorial activities of the three TFs dictate metabolic status, and that direct reduction of *Pnt* activity is sufficient to promote longevity. The role of ETS factors in longevity was not limited to *Pnt* and *Aop*. Knockdown of *Ets21c* or *Eip74EF* in distinct cell types also extended lifespan, revealing that lifespan is limited by transcription from the ETS binding site in multiple cellular contexts. Reducing the activity of the *C. elegans* ETS TF *Lin-1* also extended lifespan, a finding that corroborates established evidence of roles of this TF family in ageing. Altogether, these results reveal the ETS family of TFs as pervasive and evolutionarily conserved brokers of longevity.

## INTRODUCTION

Ageing is characterised by a steady systematic decline in biological function and increased likelihood of disease. Understanding the basic biology of ageing therefore promises to help alleviate the personal and societal burdens resulting from the increasing proportion of older people in our populations. The pursuit of this goal has revealed that ageing is plastic, and healthy lifespan can be extended by manipulating specific genes, including those encoding components of nutrient signalling pathways^1^. Such interventions often act through changes in gene expression, with transcriptional regulation playing a critical role^2–5^. Sequence-specific transcription factors (TFs) are the primary coordinators of transcriptional programmes^6^. Hence, understanding their function in adult animals will provide insight into how gene expression can be altered to optimise physiology towards promoting lasting health into late life.

TFs can be classified into families based on their DNA-binding domains, reflecting common evolutionary ancestry ^7^. TFs of the forkhead family have been studied extensively in the context of ageing. This large family of eukaryotic TFs can be further subdivided based on the sequence of their DNA-binding domain (DBD), the forkhead box. Activation of Forkhead Box O (*Foxo*) othologues in insects and nematodes extends their lifespan, and alleles of *Foxo3* are associated with longevity in humans^8–11^. Furthermore, Foxos are required for the longevity achieved by the inhibition of the insulin/IGF signalling (IIS) pathway in worms and flies^12,13^. Foxos do not act in isolation, rather their outputs are tuned by the activities of additional TFs. For example, the *C. elegans* Foxo, DAF-16, acts in concert with the heatshock factor *HSF* and the GATA family TF *Elt-2* to regulate a pro-longevity transcriptional programme^4,14^. Other TFs are regulated by IIS in parallel to DAF-16, such as SKN-1, whose activity is sufficient to extend lifespan independently of DAF-16 ^15^. The complex interactions observed between Foxos and these additional TFs are best described as regulatory circuits which must be correctly coordinated to realise anti-ageing transcriptional programmes.

The E twenty six (ETS) family of TFs, is conserved across animals, including 28 genes in humans^16–18^. The shared, defining feature of ETS TFs is a core helix-turn-helix DBD, which binds DNA on 5’-GGA(A/T)-3’ ETS-binding motifs (EBMs). These TFs are further classified into four subgroups based on variation in peripheral amino acid residues, which confer binding specificity depending on nucleotide variation flanking the core EBM. This binding specificity is thought to differentiate the transcriptional outputs of distinct ETS TFs within the same cell^19^. In common with Foxos, ETS TFs generally function as transcriptional activators, although a few have been shown to repress transcription ^20,21^. The *Drosophila Aop* (a.k.a. *Yan*, the orthologue of human *Tel*) is an ETS TF that is thought to only act as a transcriptional repressor.

The ETS family has been studied extensively in the context of cancer^18,19^, but recent evidence suggests a role in ageing. For example, AOP activation is associated with *Foxo*-mediated longevity in the fruit fly, and AOP activation alone is sufficient to extend lifespan^3^. Additionally, in multiple organisms, examination of the evolutionarily-conserved targets of Foxos has highlighted the conserved presence of ETS-binding sites within regions bound by Foxo orthologues^22^, indicating that ETS factors may be important participants in Foxo’s longevity programme in a number of animals. Accordingly, in nematodes, reducing the activity of *ets-4* extends lifespan, conditional on the presence of DAF-16^23^. Clearly, wider investigation of how ETS TFs determine lifespan is warranted.

This study set out to investigate transcriptional regulation by AOP. The study showed that *Aop* and *Foxo* drive a common longevity transcriptional programme *in vivo* and interact to determine transcriptional outcomes, both *in vitro* and *in vivo*. Additionally, *Aop* blocks transcriptional activation by *Pnt*, an ETS transcriptional activator, and *in vivo* analysis of interactions between *Aop, Pnt* and *Foxo* indicated that the key function of the common and coordinated transcriptional programme of *Foxo* and *Aop* for longevity is to block the activity of *Pnt*. The transcriptomic analysis revealed, and further experiments confirmed, that metabolism is modulated by the combined activity of the three TFs and that directly limiting *Pnt* activity was sufficient to extend lifespan. These lifespan-regulatory effects were not confined to *Pnt* and *Aop*: limiting the activities of further two ETS TFs in a range of tissues promoted longevity. The finding of a role in ageing which is conserved in at least half of the ETS TFs in *Drosophila* suggests an evolutionarily ancient origin. Consistent with this hypothesis, a previously unappreciated role in *C. elegans* longevity is shown for the ETS TF *Lin-1*. Altogether these results reveal new functions of ETS TFs at the nexus of lifespan and metabolism, opening up the investigation of these TF in adult physiology.

## RESULTS

### 1) FOXO and AOP collaboratively establish a pro-longevity transcriptional programmes

*Aop* and its orthologue *Tel* are proposed to repress transcription by physical competition with activators for binding sites^20,24,25^, recruitment of additional repressive complexes^21,25,26^ and formation of homo-oligomers to limit activator access to euchromatin^27–29^. Hence, to understand the role of *Aop* in longevity, its interactions with relevant transcriptional activators need to be examined.

Foxo is one such transcriptional activator: both *Foxo* and *Aop* are required for longevity by IIS inhibition, acting downstream of Pi3K-Akt or Ras-ERK pathways, respectively^13,23^, where the shared genomic locations bound by AOP and FOXO *in vivo* suggest that gene expression downstream of IIS is coordinated by the orchestrated recruitment of FOXO and AOP to promoters. But what is the overall relationship between the transcriptional programmes triggered by *Foxo* and *Aop*? The transcriptomic response to induction of either *Foxo*, *Aop^ACT^* (encoding a constitutively active form of AOP) or both, was characterised in adult female fly guts and fat bodies (equivalent to mammalian liver and adipose), tissues from which the two TFs extend lifespan^3^. The TF expression was under the control of the *S_1_106* driver, induced by feeding the RU_486_ inducer.

The transcriptional programmes triggered by *Foxo* or *Aop^ACT^* were significantly correlated within the set of 896 genes differentially regulated by either TF in the gut, or the equivalent 745 genes in the fat body (Figure 1A-B; for details of differential expression analysis results see Supplementary Tables 1-8). Gene Ontology (GO) enrichment analysis suggested that, in the gut, these shared targets were involved in translation and energy metabolism (Supplementary Table 9), whilst the equivalent analysis in the fat body suggested orchestration of genome regulation (Supplementary Table 10). Since the sets of differentially expressed genes were largely tissue-specific (Supplementary Figure 1), this correlated response appeared as a general feature of the *Foxo* and *Aop* regulons, independent of specific target promoters, in both the gut and fat body. Hence, *Aop* and *Foxo* appear to act on lifespan though a shared transcriptional programme.

**Figure 1.**
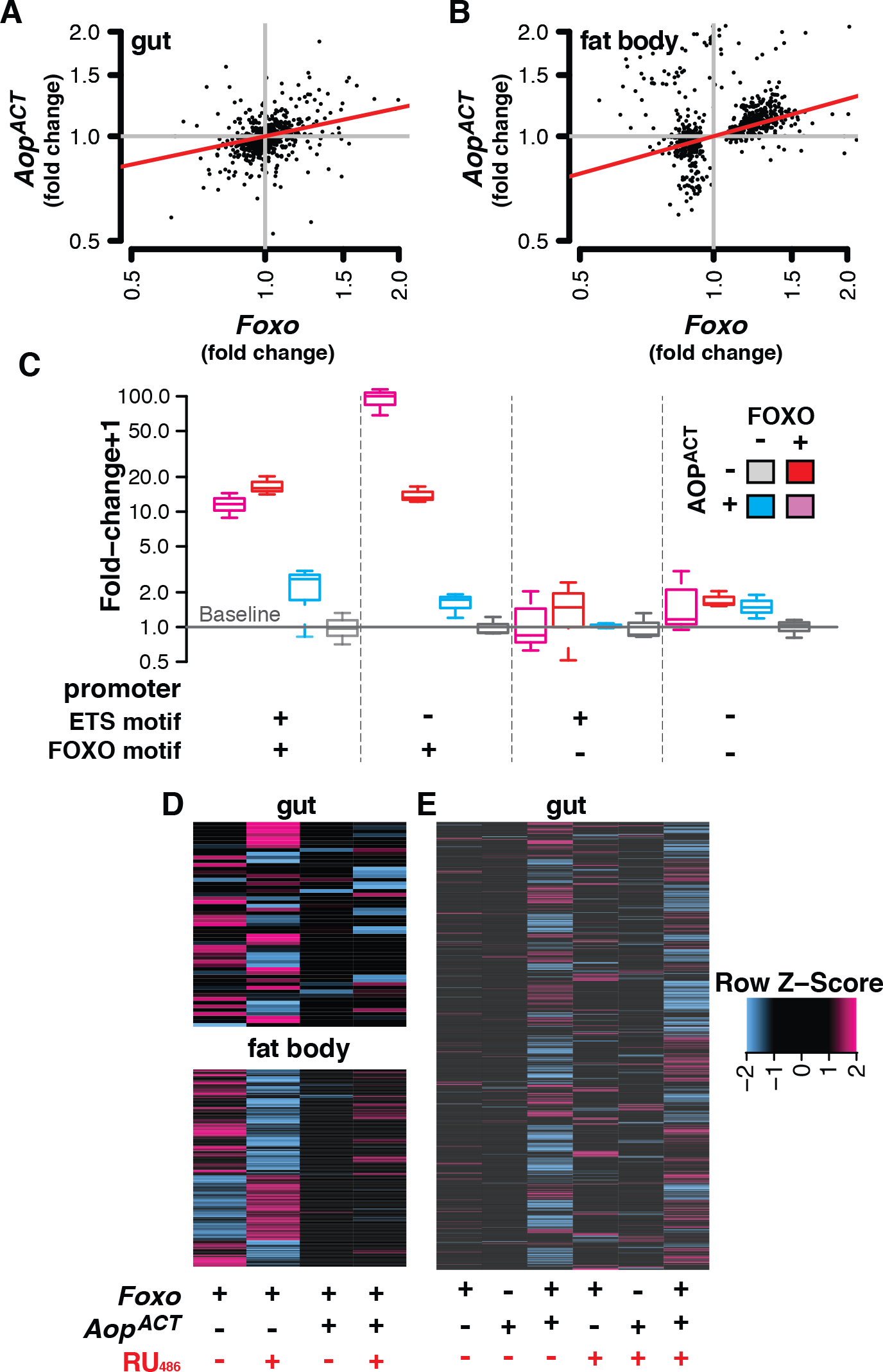
*Aop* recapitulates *Foxo*’s transcriptional output and modulates *Foxo* activity. Transcriptomic effects of *AopACT* expression in **(A)** fly gut and **(B)** fat body correlate those of *Foxo* expression. RU_486_-induced fold-changes in transcript abundance (relative to controls in the absence in RU_486_) are shown, from the union of sets of genes differentially expressed in response to either or both TFs. Red lines show correlation coefficients (Kendall’s Tau, P≤2.2e-16 for both tissues). **(C)** AOP^ACT^ both moderates and synergises transcriptional activation by FOXO on synthetic promoters containing combined ETS-binding motifs (EBMs) and FOXO-responsive elements (FREs), upstream of a basal *Adh-Firefly^luciferase^* reporter. Activity is shown following normalisation to internal *Renilla^luciferase^* controls, and calculation of fold-change over the median expression of each reporter in the absence of FOXO and AOP^ACT^. See Supplementary Figure 2 for replicate experiment. *In vivo*, co-expressing *Aop^ACT^* both **(D)** moderates the transcriptomic activity of *Foxo* in the fat body and gut, and **(E)** synergistically co-regulates transcription of distinct targets when co-expressed with *Foxo* in the gut. Values are shown as row-scaled Z values.

FOXO and AOP exhibit extensive genomic co-localisation, with 60% of FOXO-bound loci also bound by AOP in the adult female fly gut and fat body^3^. Does AOP directly modulate FOXO activity? To investigate how AOP interacts with FOXO on a promoter to influence transcription, a series of transcriptional reporters was constructed by combining the *Adh* basal promoter with FOXO-responsive elements (FREs: AACA), ETS-binding motifs (EBMs: GGAA) or both, and examined for their response to FOXO and AOP^ACT^ in *Drosophila* S2 cells (Figure 1C). Neither FOXO nor AOP^ACT^ influenced reporter expression in the absence of FREs, and FOXO alone was sufficient to activate transcription from the FREs, confirming published observations^24,30,31^. Combining the FREs and EBMs allowed AOP^ACT^ to attenuate the activation by FOXO, revealing that AOP can moderate FOXO’s activity when brought onto the same promoter. By striking contrast, in the absence of EBMs, AOP^ACT^ synergised with FOXO to stimulate induction from 20-fold by FOXO alone to 100-fold, indicating that AOP^ACT^ can accentuate FOXO’s ability to activate transcription. Since this synergy occurred in the absence of EBMs, this effect is most likely indirect. Interestingly, this synergy may account in part for the strong similarity in AOP’s and FOXO’s transcriptional programmes *in vivo*. Statistical modelling confirmed that the outcome of combining AOP and FOXO was promoter-dependent (Supplementary Table 11). Hence, the presence or absence of EBMs determines whether AOP functions to enhance or moderate FOXO activity on a promoter.

To examine if synergy and antagonism of *Foxo* by *Aop* can be observed on native promoters *in vivo*, *Foxo* targets were tested for patterns of transcriptional alteration by co-induction of *Aop^ACT^*, in the above-described transcriptomic dataset. Overall, 55 were identified in the gut (Supplementary Table 12), and 179 in the fat body (Supplementary Table 13), whose modulation by *Foxo* was attenuated by *Aop^ACT^* (Figure 1C). To determine the likely physiological outcomes of this inhibition of *Foxo* targets by *Aop*, GO enrichment analysis was performed, revealing functions in lipid catabolism in the gut (Supplementary Table 14), and genome regulation in the fat body (Supplementary Table 15). No synergistic effects could be detected in the fat body, but they could be discerned in the gut, where co-expressing both *Foxo* and *Aop^ACT^* led to differential expression of 1022 genes (Supplementary Table 16), which was not evident when either TF was overexpressed alone (Figure 1E). GO enrichment analysis suggested that these synergistically-regulated genes function in translation, proteolysis and mitochrondrial regulation (Supplementary Table 17). Thus, transcript profiling confirmed that the two modes of AOP-FOXO interaction observed on synthetic reporters can also occur *in vivo*. This simultaneous synergy and antagonism of AOP and FOXO may explain why, while activation of each TF is sufficient to promote longevity, their co-activation does not result in additive effect on lifespan^3^.

### 2) *Pnt* modulates metabolism and limits lifespan

Whilst interactions with FOXO likely account for some of the transcriptional outputs of AOP, 80% of AOP-bound sites do not appear bound by FOXO *in vivo*^3^. Given the evidence that AOP alone is insufficient to regulate transcription when brought onto a promoter (Figure 1C, and references^20,23–29^, this observation suggested that interactions with other TFs must account for the full breadth of AOP’s physiological and transcriptomic effects. One such TF is *Pnt*, whose activity is inhibited by *Aop* during fly development, which is presumed to occur by competition for binding sites since the two recognise the same DNA sequence^24,32–34^. As previously reported^24,35,36^, transcriptional induction by PNT^P1^ (a constitutively active isoform of Pnt; references^37–39^) was completely blocked by AOP^ACT^ (Figure 2A), suggesting that PNT inhibition may be a key factor in *Aop*’s pro-longevity effect. To evaluate this possibility *in vivo*, the transcriptome-wide effects of co-expressing *Aop*^*ACT*^, *Pnt*^*P1*^ *and Foxo* in the gut and fat body were assessed. In the gut, 512 transcripts appeared to be subject to the combinatorial, interactive effects of the three TFs, as were 622 in the fat body (Supplementary Table 18-19, with genes regulated by over-expressing *Pnt^P1^* alone in Supplementary Table 20-21). To reveal emergent transcriptional programmes in each tissue, principal component analysis (PCA) was performed over the transcripts that were differentially regulated by the varying combinations of the three TFs. Remarkably, the first principal component (PC) of differentially expressed genes in the gut distinguished flies by published lifespan outcomes^3^ with short-lived flies expressing *Pnt^P1^* alone or in combination with *Foxo* at one end of the PC; long-lived flies expressing one or both *Foxo* and *Aop^ACT^* forming a distinct group at the other end of the PC; and *Aop^ACT^* countering the effect of *Pnt^P1^* to form an intermediate group (Figure 2B). In the fat body, a similar grouping was apparent on the diagonal of PCs 1 and 2 (Figure 2C). To infer functional consequences of these distinct transcriptional programmes, transcripts from the input set corresponding to the PCs were isolated and GO enrichment analysis performed (Supplementary Tables 22-23; along with GO enrichment in full sets of differentially-expressed genes: Supplementary Tables 24-25). This revealed a strong enrichment of genes with roles in energy metabolism, whose expression was strongly correlated to the PCs (Supplementary Figure 3). Overall, a combined view of the PCA and GO analysis predicted that: (1) the *Foxo-Aop-Pnt* circuit regulates metabolism, and (2) inhibiting *Pnt*’s output promotes longevity.

**Figure 2.**
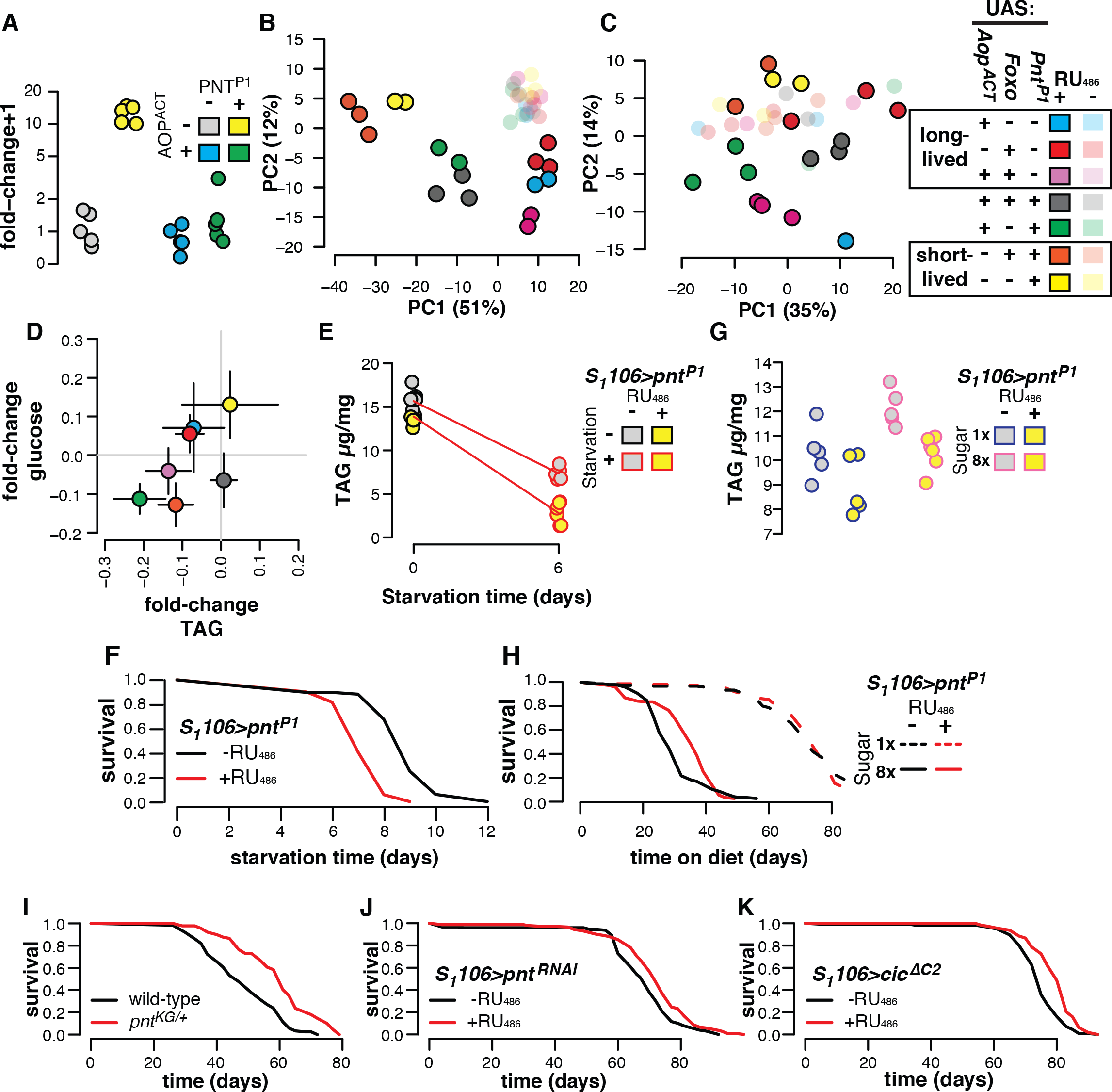
*Pnt* brokers transcriptomic, metabolic and longevity outcomes. **(A)** AOP^ACT^ counteracts activation by PNT^P1^ of a synthetic promoter containing ETS-binding motifs, upstream of an *Adh-Firefly^luciferase^* reporter. Activity is shown following normalisation to internal *TK-Renilla^luciferase^* controls, and calculation of fold-change over median expression in the absence of PNT^P1^ and AOP^ACT^. Promoter activation was subject to a significant AOPACT:PNTP1 interaction (ANOVA F_1,16_=41.725, p=7.9e-6) **(B-C)** Aggregate transcriptional effects of *Aop^ACT^* counter those of *Pnt^P1^* to establish transcriptional programs corresponding to lifespan. For gut and fat body, plots show coordinates of samples on the first two dimensions of principle components analysis, amongst transcripts which were differentially expressed according to combined TF co-expression (significant genotype:RU_486_ interaction). Legend shows samples’ groupings by previously-published lifespan outcomes resulting from TF induction in the gut and fat body or the gut alone (noting that lifespan effects of combined *Aop^ACT^* and *Pnt^P1^* expression are not known)^3^. **(D)** Metabolites are determined by a three-way interaction of *Pnt, Aop* and *Foxo*. Axes show glucose and TAG per unit body mass, expressed as fold-change (mean±SE) in the presence of RU_486_, relative to the corresponding RU_486_-negative genotype control. Both TAG and glucose were subject to statistically significant interactions of *Foxo*Aop^ACT^*Pnt^P1^* induction (Supplementary Tables 20-21). **(E)** Over-expressing *Pnt^P1^* in the gut and fat body accelerates loss of TAG under starvation stress. (ANOVA RU_486_:starvation F_1,19_=7.03, p=0.02. Full statistical analysis in Supplementary Table 30). **(F)** Over-expressing *Pnt^P1^* in the gut and fat body reduces survival under starvation stress. The plot shows 71 deaths with RU_486_, 68 deaths without, p=1.3E-14 (log-rank test). **(G)** Over-expressing *Pnt^P1^* in the gut and fat body reduces accumulation of TAG on a high-sugar diet. *Pnt^P1^* reduced TAG (ANOVA F_1,17_=14.4, p=1.4e-3), sugar increased TAG (F_1,17_=15.25, p=1.1e-3), with *PntP1* appearing to counteract the effect of sugar (mean±SE: control 10.31±0.48, high sugar * RU_486_ 10.30±0.29; t-test t=0.009, p=0.99, Full statistical analysis in Supplementary Table 31). **(H)** Over-expressing *Pnt^P1^* in the gut and fat body enhances survival on a high-sugar diet. Plot shows 139 deaths and 3 censors on high sugar with RU_486_ feeding (median=33.5 days), 146 deaths and 1 censor on high sugar without RU_486_ feeding (median=26.5 days), 133 deaths and 17 censors on low sugar with RU_486_ feeding (median=73 days), 122 deaths and 28 censors on low sugar without RU_486_ feeding (median=71 days; Cox Proportional Hazards RU_486_:diet p=6e-3; Full statistical analysis in Supplementary Table 32). **(I)** Heterozygous *Pnt* mutants are long-lived. Plot shows 122 wild-type deaths (median=45.5 days), 89 *pnt^KG^/+* deaths (median=59.5 days), log-rank test p=9.2e-11. **(J)** Adult-onset *Pnt* inhibition in the gut and fat body is sufficient to extend lifespan. Plot shows 132 deaths and 19 censors without RU_486_ feeding (median=68.5 days), 139 deaths and 16 censors with RU_486_ feeding (median=71 days), log-rank test p=7.2e-4. **(K)**Overexpression of *Cic*, a transcriptional repressor of *Pnt*, is sufficient to extend lifespan. Plot shows 113 deaths and 9 censors without RU_486_ feeding (median=73.5 days), 113 deaths and 8 censors with RU_486_ feeding (median=78.5 days), log-rank test p=1.5e-7.

To test the prediction that the *Foxo-Aop-Pnt* circuit regulates metabolism, the individual and combined effects of the three TFs were tested on protein, TAG (triacylglyceride, the main energy store in insects) and glucose. Feeding RU_486_ to *S106* control flies did not affect TAG, protein or glucose content (Supplementary Figure 4). Since body mass was subject to a complex interaction involving all three TFs (Supplementary Figure 5, Supplementary Table 26), confounding per-fly quantification, protein, glucose and TAG were normalised to body mass. Protein density was increased by *Foxo* overexpression, and this effect was enhanced by *Pnt* co-expression (Supplementary Figure 6, Supplementary Table 27). *Foxo* and *Aop^ACT^* reduced TAG, but had no effect on glucose (Figure 2D). By contrast, flies over-expressing *Pnt^P1^* had moderately increased whole-body glucose, with no evidence of alterations to TAG in this experiment (Figure 2D). Critically, the metabolic effects of each TF were highly dependent on the activities of the others for both glucose and TAG levels, as well as overall body mass, which was confirmed with statistical analyses (Supplementary Tables26 and 28-29). Overall, metabolite profiling revealed a tripartite dialogue between the three TFs, in which distinct combinations have unique outcomes on metabolism, hence confirming the physiological prediction from the transcriptomic analysis.

Since *Pnt* appeared to dictate the transcriptional outcomes that predicted metabolic regulation by the *Foxo-Aop-Pnt* circuit, the role of *Pnt* in responses to nutritional stress was further evaluated. TAG was quantified after a week of *Pnt* over-expression, and then after a subsequent six days of starvation. *Pnt^P1^* over-expression increased the loss of TAG induced by starvation (Figure 2E), suggesting that PNT activation predisposes flies to mobilise energy stores. This was associated with decreased resistance to the starvation stress, with flies over-expressing Pnt dying 24% earlier on average (Figure 2F). The observed ability of *Pnt* to promote catabolism of energy stores may be beneficial in the face of over-nutrition, and relevant to the Western human epidemic of metabolic disease associated with energy-rich diets. A *Drosophila* model of such energy-rich diets is increasing dietary sugar. Flies fed a 40% sugar diet die substantially earlier than controls fed a 5% sugar diet, and accumulate TAG ^40–42^. However *Pnt^P1^* overexpression restored TAG levels in flies on a high-sugar diet to those observed on a low-sugar diet (Figure 2G). Whilst there was no statistically significant interaction of sugar and *Pnt^P1^* induction in a linear model (Supplementary Table 31), the adipogenic effect of sugar was opposed by *Pnt*, such that TAG levels on a high sugar-diet with RU_486_ were equivalent to those on a low-sugar diet without RU_486_ (t-test: t=0.01, p=0.99). Moreover, *Pnt^P1^* induction spared flies from the full extent of the early death induced by dietary sugar, increasing median survival time by 26%, despite having no effect on the low-sugar diet (Figure 2H). Note that in two of three experiments performed, and consistent with published data^3^, the induction of *Pnt^P1^* reduced the basal levels of TAG (Figure 2E and 2G). Altogether, these results suggest that complex interactions in the *Foxo-Aop-Pnt* circuit determine homeostatic set-points for nutrient storage; and that *Pnt* predisposes flies to leanness, which correlates survival of nutritional stress.

But what is the role of *Pnt* under healthy, nutritionally-optimal conditions? The attenuation of *Pnt*’s transcriptomic effects by *Aop* indicated that limiting *Pnt* activity directly may be sufficient to extend lifespan. To directly reduce *Pnt*, a validated^43^ loss-of-function p-element insertion in *Pnt* (*Pnt^KG04968^*, henceforth *Pnt^KG^*), was backcrossed into an outbred, wild-type background for ten generations. The mutation was homozygous lethal. However, heterozygote females exhibited a 20% increase in median lifespan (Figure 2I). Similarly, inducing RNAi against *Pnt* from day three of adulthood in the gut and fat body also increased lifespan (Figure 2J). The HMG-box repressor *capicua* (*cic*) represses expression of ETS factors including *Pnt* ^44^ and, consistent with the effects of *pnt^RNAi^*, overexpressing *cic^**∆**C2^* (a *cic* mutant lacking a known MAPK phosphorylation site) in the gut and fat body also substantially extended lifespan (Figure 2K). This further confirmed the functional predictions from the transcriptomic analysis and demonstrated that countering *Pnt*, in the tissues in which *Foxo* and *Aop* over-expression is beneficial, is sufficient to extend lifespan.

### 3) Modulating the activity of multiple ETS factors in multiple tissues and distinct species extends lifespan

Animal genomes encode multiple ETS factors. In *Drosophila* the ETS family comprises *Aop* and *Pnt* along with six other ETS TFs (*Ets21c, Ets65A, Eip74EF, Ets96B, Ets97D, Ets98B*), each of which is expressed with its own unique tissue-specific pattern (Supplementary Figure 7). The presumed common ancestry of these TFs, along with their tissue-specific expression, suggests that they may regulate common functions in distinct or partially-overlapping tissues, in which case roles in longevity may extend beyond *Aop* and *Pnt*. *Cic*, whose overexpression extended lifespan (Figure 2K), represses both *Pnt* and *Ets21C* ^44^, suggesting that *Ets21C* may also limit lifespan. Similar to *pnt^RNAi^* and *cic^**∆**C2^*, inducing *Ets21C^RNAi^* in the gut and fat body with *S_1_106* extended lifespan (Figure 3A). Furthermore, both heterozygous and homozygous mutants of *Ets21C* (bearing *Ets21C^f03639^*, henceforth Ets21C^F^, an intronic P-element insertion which was backcrossed 10 times into wild-type flies) were also long-lived relative to controls (Figure 3B). Thus, lifespan limitation is conserved between *Pnt* and *Ets21c*.

**Figure 3.**
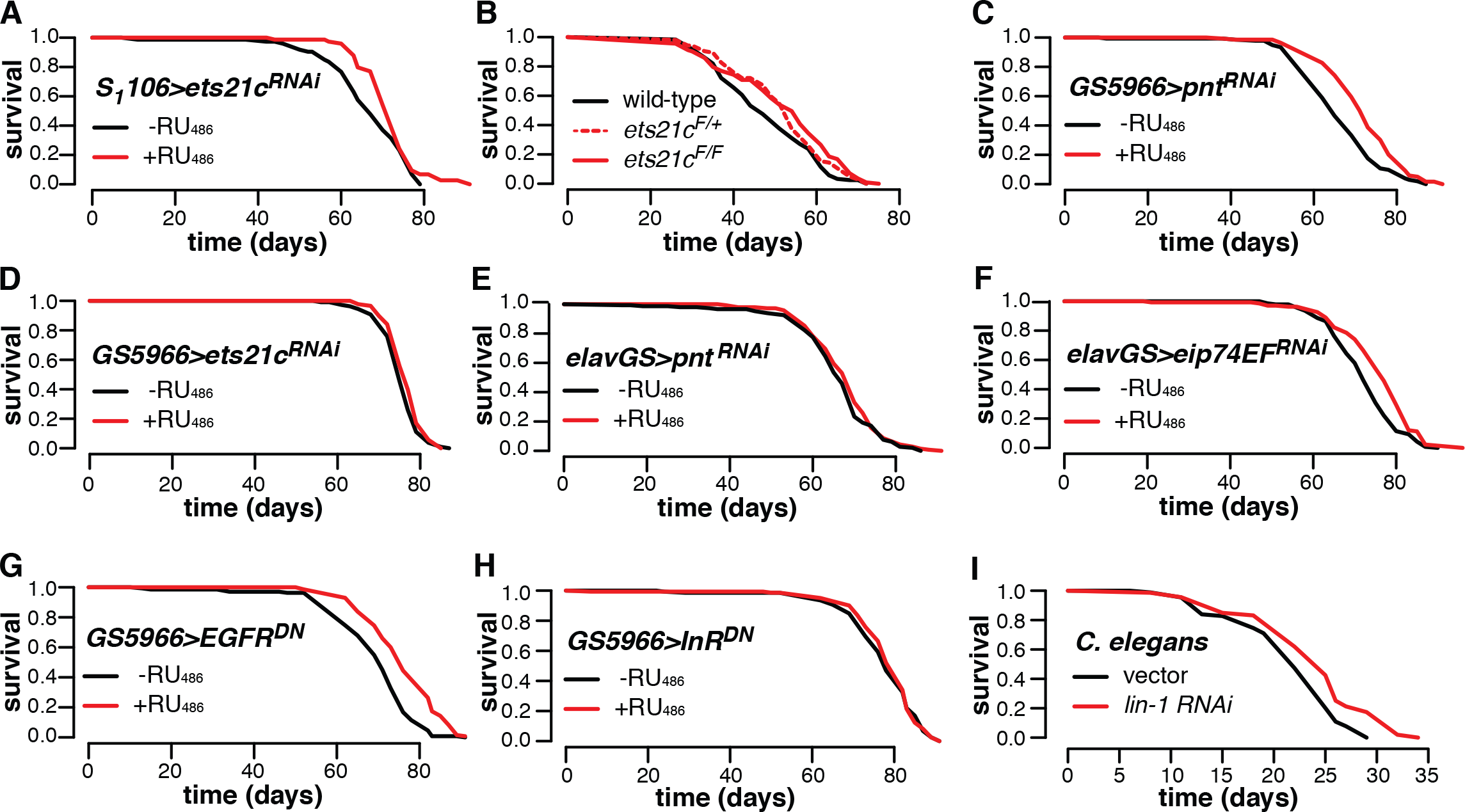
ETS transcription factors limit lifespan amongst diverse *Drosophila* tissues and across species. **(A)** Knockdown of Ets21c by expressing *Ets21c^RNAi^* in the gut and fat body extends lifespan. Plot shows 72 deaths and 1 censor without RU_486_ feeding (median=65.5), 74 deaths and 1 censor with RU_486_ feeding (median=71), p=0.05 (log-rank test). **(B)** Systematic *Ets21c* knockdown extends lifespan. Plot shows 122 wild-type deaths (median=45.5), 113 *Ets21c^F^/+* deaths (median=52.5) and 104 *Ets21c^F^*/*Ets21c^F^* deaths (median=52.5). Heterozygous p=0.05, homozygous p=0.003 (log-rank tests). Note that wild-type controls are also presented in Figure 2I. **(C)** Expressing *Pnt^RNAi^* in enterocytes extends lifespan. Plot shows 121 deaths and 29 censors with RU_486_ feeding (median=63.5), 127 deaths and 23 censors without RU_486_ feeding (median=72), p=7.33e-6 (log-rank test). **(D)** Expressing *Ets21c^RNAi^* in enterocytes does not affect lifespan. Plot shows 108 deaths and 12 censors without RU_486_ feeding (median=73.5), 88 deaths and 17 censors with RU_486_ feeding (median=76), p=0.07 (log-rank test). **(E)** Expressing *PntRNAi* in neurons did not affect lifespan. Plot shows 140 deaths and 10 censors without RU_486_ feeding (median=68.5), 146 deaths and 4 censors with RU_486_ feeding (median=66), p=0.27 (log-rank test). **(F)** Expressing *Eip74EFRNAi* in neurons extends lifespan. Plot shows 140 deaths (and no censors) without RU_486_ feeding (median=71), 134 deaths and 6 censors with RU_486_ feeding (median=76), p=6.73e-5 (log-rank test). **(G)** Expressing a dominant-negative form of *EGFR* (*EGFRDN*) in enterocytes extends lifespan. Plot shows 132 deaths and 16 censors without RU_486_ feeding (median=70), 135 deaths and 15 censors with RU_486_ feeding (median=74.5), p=2.2e-9 (log-rank test). **(H)** Expressing a dominant-negative form of *InR* (*EGFR^DN^*) in enterocytes does not affect lifespan. Plot shows 127 deaths and 23 censors without RU_486_ feeding (median=77), 134 deaths and 16 censors with RU_486_ feeding (median=80), p=0.66 (log-rank test). **(I)** Knocking down *lin-1* in the nematode *C. elegans* extends lifespan. Plot shows 83 deaths and 6 censors with *lin-1^RNAi^* feeding (median=25) and 100 deaths and 4 censors in vector-fed controls (median=22), p=3.3e-2.

Are *Ets21c* and *Pnt* relevant to lifespan in the same tissues? To target a subset of the gut and fat body cells marked by *S_1_106*, both TFs were knocked down specifically in enterocytes using the inducible, enterocyte-specific driver *GS5966*. *Pnt* knockdown in enterocytes alone was still sufficient to extend lifespan (Figure 3C). However, expressing *Ets21c^RNAi^* with the same driver had no effect on longevity in these cells (Figure 3D). This specificity appeared to reflect tissue-specific lifespan-limiting function of *Pnt*, rather than differences in expression, since *Ets21c* is more highly expressed in these cells than *Pnt* ^45^, therefore suggesting a level of tissue specificity in ETS TFs’ effects on ageing. In this case, knocking down diverse ETS factors in diverse tissues might be expected to extend lifespan.

*Pnt* is of known relevance to neurophysiology ^46^, especially in neurogenesis, and its continued expression in adults^5^ suggests an ongoing physiologically-relevant role in neurons. However, expressing *Pnt^RNAi^* in neurons using the *Elav-GS* driver did not affect lifespan (Figure 3E). To explore if other ETS TFs might be relevant to lifespan in neurons, *Eip74EF* was targeted by neuronal RNAi, because it is more highly expressed in adult brain than in any other tissue (Supplementary Figure 7). We found that this intervention also extended lifespan (Figure 3F). Hence, the *Drosophila* ETS family includes at least four TFs with roles in ageing (*Aop, Pnt, Ets21C, Eip74EF*), and with distinct lifespan-limiting effects in specific tissues.

The ETS TFs act downstream of receptor tyrosine kinase pathways^13,44^. We also found some evidence that different RTKs limit longevity in different cells: inducing the dominant-negative form of the epidermal growth factor receptor (EGFR^DN^) in enterocytes extended lifespan (Figure 3G), phenocopying knockdown of *Pnt*, whereas the induction of the dominant-negative insulin receptor (InR^DN^) did not (Figure 3H), whilst it is known to extend lifespan under control of other drivers^10^, and even though it is the more highly expressed of the two RTKs in these cells^45^. Hence, different ETS factors may limit lifespan downstream of different RTK pathways.

Overall, we found evidence that the role in ageing is shared by multiple ETS factors in *Drosophila*. ETS TFs are conserved throughout multicellular animals. The genome of the nematode *Caenorhabditis elegans* encodes 11 in total. At least one of these, ETS-4, has been reported to limit lifespan in the worm intestine^23^. We found that the knockdown of one more, *Lin-1*, can also extend *C. elegans* lifespan (Figure 3I). Thus, multiple ETS factors limit lifespan across hundreds of millions of years of evolutionary divergence, hinting at a general role for this family of TFs across animals.

## DISCUSSION

Promoting lifespan by transcriptional control is an attractive prospect, because targeting one specific protein can restructure global gene expression to reprogram physiology, orchestrating a cellular recalibration that provides broad-scale benefits in ageing. This study suggests key roles for ETS TFs in such optimisation. The results show dual roles for the ETS repressor *Aop* in longevity: balancing *Foxo*’s outputs, whilst also opposing *Pnt*’s outputs. The outcome of this tripartite interaction is a program of expression that corresponds to lifespan including genes with critical and conserved roles in central metabolism. That directly reducing physiological *Pnt* is sufficient to extend lifespan suggests that repressing transcription from the ETS site is the key longevity-promoting step in this circuit. Accordingly, reducing transcription from the ETS site by targeting multiple TFs, in multiple *Drosophila* tissues, and in multiple animal taxa, was sufficient to extend lifespan. Altogether, these results show that inhibiting lifespan is a general feature of ETS transcriptional activators, and furthermore that this feature is conserved across the ETS family.

The apparent lifespan-limiting role of ETS factors in adult animals is doubtlessly balanced by selection for their important roles in development^24^. There are at least two possible explanations for why these TFs with detrimental long-term effects are active in adult tissues. ETS TF activity may simply run-on from development, which may be neutral during the reproductive period (i.e. when exposed to selection) but costly in the long term in aged animals. Additionally or alternatively, there may be context-dependent benefits of activating transcription from the ETS site. This latter explanation is supported by two lines of evidence: the enrichment of metabolic functions in the *Pnt* and *Aop* regulons suggests that metabolic homeostasis is determined not by either TF alone, but by the balance of activation versus repression of transcription from the ETS site, in which case benefits of *Pnt* activity would only be exposed in the face of metabolic variation or stress. Fully consistent with the notion of context-dependent benefits of activating transcription from the ETS site, the present data show that whilst *Pnt* is costly on a low-sugar diet, it can improve survival on a high-sugar diet. The second line of evidence for context-dependent benefits of *Pnt* is that, in mammals, different ETS TFs have distinct but partially-overlapping binding profiles^16,19^, which may indicate a shared set of core functions - perhaps in metabolism - which are important in response to distinct signalling cues. In this case, the outcome of activating an ETS TF would depend on the fit between promoter architecture and domains on the TF protein adjacent to the ETS domain. Indeed, the unexpected recent finding that neuronal *Pnt* and *Aop* can have positively-correlated transcriptomic effects^47^ is consistent with highly context-dependent ETS TF function, likely subject to complex interactions of euchromatin availability, a given TF’s complement of protein domains, and the status of intracellular signalling networks. This context-dependence makes it all the more remarkable that roles in lifespan appear to be a conserved feature of the ETS TFs in diverse contexts.

Tissue environment appears to be a key contextual factor determining the effects of ETS TFs on lifespan. This study shows that inhibiting distinct ETS TFs in multiple tissues is sufficient to extend lifespan. Differences between tissues in chromatin architecture are likely to alter the capacity of a given ETS TF to bind a given site. Indeed, whilst *Pnt* is of known neurophysiological importance, the lifespan extension mediated by expressing RNAi against *Pnt* in the gut and fat body was not recapitulated by expressing the same construct in neurons. By contrast, neuronal RNAi against *Eip74EF* was sufficient to extend lifespan, and it has previously been implicated as a mediator of age-dependent functional decline^48^. Similar tissue-specific effects were evident in the matrix of *Pnt* and *Ets21c* knockdown in the gut and fat body versus the gut alone. Furthermore, overexpression of dominant-negative receptor tyrosine kinases which are known to act upstream of ETS TFs also had tissue-specific effects: over-expressing dominant-negative *EGFR^DN^* in enterocytes extended lifespan, whilst the equivalent expression of *InR^DN^* did not, despite the status of both receptors as activators of Ras/ERK signalling^13,44^. This correspondence between the tissue-specific outcomes of *Ets21c* and *InR* knockdown, and of *Pnt* and *EGFR* knockdown, suggests that lifespan-promoting transcriptional programs may be inhibited by similar cellular signalling cascades across tissues, such as *Pi3K-Akt* and *Ras-ERK*, but local regulation is coordinated by distinct receptors and TFs which nevertheless ultimately converge on the ETS binding motif. It follows to ask whether inhibiting activation from the ETS site in different tissues extends lifespan by alike cell-autonomous effects, or whether these genetic lesions lead to cell-nonautonomous effects which are common across different tissues, giving an organismal benefit that manifests in longevity. The relative importance in ageing of cell-autonomous versus nonautonomous outputs of local transcriptional control remains to be established.

The structure of molecular networks and their integration amongst tissues underpins phenotype, including into old age. This should be a key therapeutic consideration, as we attempt to bridge the gap between genotype and specific age-related pathologies, such as dementia or cancer. Thus, unravelling the basics of these networks is a critical step in identifying precise anti-ageing molecular targets^1^. Perturbing specific regulatory hubs can identify potential therapeutic targets, and identifying the least disruptive perturbation of these networks, by targeting the “correct” effector, is a key goal in order to achieve desirable outcomes without undesirable tradeoffs that may ensue from broader-scale perturbation. This targeting can be at the level of specific proteins, specific cell types, specific points in the lifecourse, or a combination of all three. The tissue-specific expression pattern of ETS TFs, and the apparent conservation of their roles in longevity across distinct tissues, ETS family members and animal phyla, highlights them as important regulators of tissue-specific programs, which may be beneficial in medically targeting both lifespan and precise senescent pathologies.

## MATERIALS & METHODS

### *D.melanogaster* culture

All experiments were carried out in outbred, *Wolbachia*-free *Dahomey* flies, bearing the *w1118* mutation and maintained at large population size since original domestication. All transgenes (Supplementary Table 33) were backcrossed into this background at least 6 times prior to experimentation and maintained without bottlenecking. Cultures were maintained on 10% yeast (MP Biomedicals, OH, USA), 5% sucrose (Tate & Lyle, UK), 1.5% agar (Sigma-Aldrich, Dorset, UK), 3% nipagin (Chemlink Specialities, Dorset, UK), and 3% propionic acid (Sigma-Aldrich, Dorset, UK), at a constant 25°C and 60% humidity, on a 12:12 light cycle. Experimental flies were collected as embryos following 18h egg laying on grape juice agar, cultured at standardised density until adulthood, and allowed to mate for 48h before males were discarded and females assigned to experimental treatments at a density of 15 females/vial. To induce transgene expression using the GeneSwitch system, the inducer RU_486_ (Sigma M8046) was dissolved in absolute ethanol and added to the base medium to a final concentration of 200 *μ*M. Ethanol was added as a vehicle control to RU-negative food. For lifespan experiments, flies were transferred to fresh food and survival was scored thrice weekly. Feeding RU_486_ to driver-only controls did not affect lifespan (Supplementary Figure 8). For starvation stress experiments, flies were fed RU_486_ or EtOH-supplemented media for one week, before switching to 1% agarose with the equivalent addition of RU_486_ or EtOH, with death scored daily until the end. For sugar stress experiments, sugar content was increased to 40% w/v sucrose^40–42^.

### *C. elegans* culture

Worms were maintained by the protocol of Brenner^49^, at 20°C on NGM plates seeded with *Escherichia coli* OP50. For lifespan experiments, N2 (wildtype N2 male stock, N2 CGCM) were used at 20°C on NGM plates supplemented with 15μM FUDR to block progeny production. RNAi treatment was started from egg. Animals that died from internal hatching were censored.

### Molecular cloning

The *pGL3Basic-4xFRE-pADH-Luc* construct (called pGL4xFRE) described in reference^31^ was used as template to generate PCR products containing 6xETS-4xFRE-pADH, 4xFRE-pADH, 6xETS-pADH- or pADH (primers in Supplementary Table 34, ETS sequence as described by reference^50^), flanked by *XhoI* and *HindIII* sites, cloned into the corresponding sites in pGL3Basic and confirmed by sequencing.

*PntP1* was amplified from *UAS-PntP1* genomic DNA with Q5 High-Fidelity Polymerase (NEB M0491S - primers in Supplementary Table 26) *Aop^ACT^* was cloned from genomic DNA of *UAS-Aop^ACT^* flies as described in ^3^. *PntP1* and *Aop^ACT^* sequences were then cloned into the *pENTR-D-TOPO* gateway vector (Thermo 450218) before recombination into the *pAW* expression vector.

### S2 cell culture

*Drosophila* S2 cells were cultured in Schneider’s medium (Gibco/Thermo Scientific 21720024), supplemented with 10% FBS (Gibco/Thermo Scientific A3160801) and Penicillin/Streptomycin (Thermo 15070063). Cells were split into fresh media 24h before transfection, then resuspended to a density of 10^6^ ml^−1^ and transfected using Effectene reagent (Qiagen 301425) in 96-well plates, according to the manufacturer’s instructions. Reporters and TF expression plasmids were co-transfected with *pAFW-eGFP* to visually confirm transfection, and *pRL-TK-Renilla^luc^* as an internal control for normalisation of reporter-produced Firefly luciferase. Reporter activity was measured 18h after transfection using Dual-Luciferase reagents (Promega E1960). *pAHW-Foxo* and/or *pAW-Aop^ACT^* were co-transfected with promoters bearing combinations of FREs and EBMs. *pAW-Aop^ACT^* and *pAW-Pnt^P1^* were co-transfected with a promoter bearing EBMs.

### Transcriptomics

Fly guts and fat bodies were dissected in ice-cold PBS and placed directly into ice-cold Trizol (Ambion 15596026), from flies bearing combinations of *UAS-Foxo*, *UAS-Aop^ACT^* and *UAS-PntP1* in an *S1_1_06-GS* background, after six days adult feeding on RU_486_. Three experimental replicates were sampled for all conditions, each comprising a pool of twelve fat bodies or guts. RNA was extracted by Trizol-chloroform extraction, quantified on a NanoDrop, and quality-assessed on an Agilent Bioanalyzer. Poly(A) RNA was pulled down using NextFlex Poly(A) beads (PerkinElmer NOVA-512981). Samples with low yields or low quality of RNA were excluded, leaving 2-3 replicate samples per experimental condition. RNA fragments were given unique molecular identifiers and libraries were prepared for sequencing using NextFlex qRNAseq v2 reagents (barcode sets C and D, PerkinElmer NOVA-5130-14 and NOVA-5130-15) and 16 cycles of PCR. Individual and pooled library quality was assessed on an Agilent Bioanalyzer. Sequencing was performed on an Illumina HiSeq 2500 instrument by the UCL Cancer Institute.

### Metabolic assays

Metabolites were measured as per reference^51^ in whole adult flies after setting up the same fly genotypes as for transcriptomics and an additional *S_1_106/+* control, and following one week of RU_486_ feeding. Flies were CO_2_-anaesthetised, weighed on a microbalance, and immediately flash-frozen in liquid N_2_. To assay metabolites, flies were thawed on ice and homogenised by shaking with glass beads (Sigma G8772) for 30s in a ribolyser at 6500 Hz in ice-cold TEt buffer (10 mM Tris, 1 mM EDTA, 0.1% v/v Triton-X-100). Aliquots of these homogenates were spun 1m at 4500g and 4°Cto pellet debris, and re-frozen at - 80°C for protein quantification. Protein was assayed with the Bio-Rad Protein DC kit (Bio-Rad 5000112). A second set of aliquots were heated to 72°C for 15m to neutralise enzymatic activity, before spinning and freezing prior to triglyceride and carbohydrate assays. Triglyceride was measured by treating 10 *μ*l sample with 200 *μ*l Glycerol Reagent (Sigma F6428) for 10m at 37°C and measuring absorbance at 540 nm, then incubating with 50 *μ*l Triglyceride Reagent (Sigma F2449) for 10m at 37°C and re-measuring absorbance at 540 nm, calculating glycerol content in each reading, then quantifying triglyceride content as the difference between the first and second measurement. Glucose was measured on 5 *μ*l homogenates with Infinity reagent (Thermo TR15421) after 15m incubation at 37°C.

### Data analysis

Sequence data were quality-checked by FastQC 0.11.3, duplicate reads were removed using Je 1.2, and reads were aligned to *D. melanogaster* genome 6.19 with HiSat2 2.1. Alignments were enumerated with featureCounts 1.6. All downstream analyses were performed in R 3.3.1. The gut and fat body were analysed in parallel. Transcripts with a mean <1 read count were excluded, leaving 11069 in the fat body and 10366 in the gut (Supplementary Tables 35-36). Read counts are given in Supplementary Tables 37-40. Differentially expressed (DE) genes were identified using DESeq2, at a false discovery rate of 10%. Effects of RU_486_ feeding were established in individual genotypes, and for specific analyses the interactive effects of genotype and RU_486_ were established. Sets of shared *Foxo* and *AopACT* targets were formed as the union of DE genes in *S106* flies over-expressing one or both transcription factors, following RU_486_ feeding. Epistatic interactions amongst TFs were identified by fitting models of the form

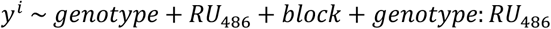

where *block* represented experimental replicate. The tripartite interaction of *Foxo*, *Aop^ACT^* and *Pnt^P1^* was identified by applying the model to all genes across all experimental conditions, and isolating genes with a significant *genotype:RU_486_* term. Antagonism of *Foxo*‘s outputs by *AopACT* was identified by fitting the model to samples of flies bearing either *UAS-Foxo* or both *UAS-Foxo* and *UAS-Aop^ACT^*, on the subset of genes which had already been identified as DE following *Foxo* over-expression. Synergistic effects of *Foxo* and *Aop^ACT^* were identified in the gut by fitting the model to all genes, from samples bearing either or both of the *UAS-Foxo* and *UAS-Aop^ACT^* transgenes. GO analysis was performed using the TopGO package, applying Fisher’s test with the *weight01* algorithm. Principal Components Analysis was performed on read counts of these genes following a variance-stabilizing transformation. To characterise gene-expression correlates of principal components, loadings onto principal components were extracted using the *dimdesc* function from the *FactoMineR* library, and GO analysis performed as previously. Transcripts of genes annotated with enriched GO terms were then plotted per term by centering variance-stabilised reads to a mean of zero and plotting against PC values per sample. Heatmaps were plotted using the *heatmap.2* function from the *gplots* library, ordering rows by hierarchical clustering by Ward’s method on Euclidian distance, and scaling to row.

Fly lifespan data were analysed using log-rank tests in Microsoft Excel. Worm lifespan data were analysed by log-rank tests in JMP. Luciferase reporter data were normalised by taking the ratio of firefly luciferase to renilla luciferase signal and, for each promoter, calculating fold-change for each sample relative to the median activity of the promoter in the absence of FOXO and AOP^ACT^. To assess the interaction of FOXO and AOP with promoters’ complements of TF-binding motifs, these normalised data were analysed by fitting a linear model of the form

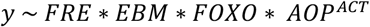

in which *y* was the *log_N_* of fold-change+1, FRE and EBM represented the TF-binding complement, and FOXO and AOP^ACT^ represented co-transfection with *pAHW-Foxo* or *pAW-Aop^ACT^*. The interactive effect of PNT^P1^ and AOP^ACT^ were assessed by fitting a linear model of the form

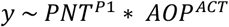

The interactive effect of TFs on metabolites and body mass were analysed by normalising metabolite density to fly mass, and then calculating fold-change for each experimental genotype in the presence of RU_486_ relative to the mean in the absence of RU_486_. These fold-change data per metabolite were analysed by linear models of the form

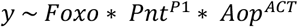

where each predictor encoded a binary term for the presence/absence of the TF. The effect of *Pnt^P1^* overexpression on TAG and lifespan responses to nutrient stress (starvation or high-sugar diet) were analysed by a model of the form

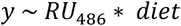

where y represented TAG normalised to unit weight in a linear model, or survival in a Cox Proportional Hazards model (survival library).

## ACKNOWLEDGMENTS

We thank Danny Filer, Rita Ibrahim, Kami Shalfrooshan, Jeremie Subrini and Caesar Chi for technical assistance; Oscar Puig, Rachel Hunt, Joeseph Bateman, Cathy Slack and Ekin Bolukbasi for sharing plasmids and cells; Bruce Edgar for sharing flies; Steve Parratt for analytical advice; and Jennifer Lohr for conducting pilot experiments in *C. elegans*. This work was funded by BBSRC grant BB/M029093/1 and Royal Society grant RG140694 to NA.

## SUPPLEMENTARY FIGURE LEGENDS

**Figure S1.** Euler plot showing overlap between the unions of *Foxo* and *Aop*’s regulons in the gut and fat body.

**Figure S2.** Replicate experiment of results shown in Fig 3c. Results were qualitatively consistent in each of the two replicates: AOP^ACT^ both moderates and synergises with transcriptional activation by FOXO on synthetic promoters containing combined ETS-binding motifs (EBMs) and FOXO-responsive elements (FREs), upstream of a basal *Adh-Firefly^luciferase^* reporter. Activity is shown following normalisation to internal *Renilla^luciferase^* controls, and calculation of fold-change over the median expression of each reporter in the absence of FOXO and AOP^ACT^. Statistical analysis in Supplementary Table 11.

**Figure S3.** Expression of transcripts subject to the 3-way *Foxo-Aop-Pnt* interaction, and annotated with significantly enriched GO terms (the five categories with lowest enrichment p-values), plotted over the first principal component of the expression matrix for each tissue (correlated to lifespan). Expression values were derived by applying *DESeq2*’s variance-stabilising transformation to read counts, taking medians per transcript, and mean-sweeping values. Principal component values are shown in Figure 2B-C.

**Figure S4.** Activating *Gal4* in the fat body and gut by feeding the inducer RU_486_ to *S_1_106/+* control flies does not affect whole-body levels of TAG (t-test t=-0.61, df=11.54, p=0.56) or glucose (t-test t=1.27, df=13.98, p=0.22). Data were collected in the same experiment as shown in Figure 2D.

**Figure S5.** Effects of *Foxo-Aop-Pnt* interactions on body mass per fly. X-axis labels indicate the combination of overexpression constructs (e.g. “foxo,aop,pnt” denotes presence of *UAS-Foxo*, *UAS-Aop^ACT^* and *UAS-PntP1*), with *S_1_106* present in all cases. “+” indicates *S_1_106*-only controls. Accompanying statistical analysis is presented in Supplementary Table 26.

**Figure S6.** Effects of *Foxo-Aop-Pnt* interactions on protein content per unit fly weight. X-axis labels indicate the combination of overexpression constructs (e.g. “foxo,aop,pnt” denotes presence of *UAS-Foxo*, *UAS-AopACT* and *UAS-PntP1*), with *S_1_106* present in all cases. “+” indicates *S_1_106*-only controls. Accompanying statistical analysis is presented in Supplementary Table 27.

**Figure S7.** Tissue-specific expression of ETS TFs. A variance-stabilising transformation was applied to read counts (from^5^, from the same population of flies, on the same media) and medians were calculated. Data were column-scaled so that the figure shows each TFs relative expression across tissues, and hierarchically clustered using Ward’s method on Euclidian distance.

**Figure S8.** Activating *Gal4* in the fat body by feeding the inducer RU_486_ to *S_1_106/+* control flies, or in enterocytes in *GS5966/+* control flies, does not affect lifespan. *GS5966/+*: plot shows 125 deaths and 4 censors (median=71) without RU_486_ feeding, 128 deaths and 6 censors with RU_486_ feeding (median=73.5), p=0.08 (log-rank test). *S_1_106/+*: plot shows 123 deaths and 3 censors without RU_486_ feeding (median=73.5), 128 deaths and 6 censors with RU_486_ feeding (median=73.5).

